# Near chromosome-level genome assembly of *Neomusotima conspurcatalis* gives insights into the evolution of moth genome architecture and fern-insect interactions

**DOI:** 10.64898/2025.12.17.694975

**Authors:** Jessie A. Pelosi, Taylor R. Curry, Abby R. Pearse, Melissa C. Smith, Katrina M. Dlugosch

## Abstract

Plant-insect interactions are the foundation of ecosystems globally, yet we are still determining the underlying mechanisms through which these relationships evolve. The co-evolution between insects and their host plants should shape the genomes of both partners, and genes involved in interaction specificity should show unique genomic signatures (e.g., rapid evolution, gene family expansions). Biological control programs are an excellent system for disentangling the genomics and molecular biology of the establishment of an insect and its host plant specificity. Fern-insect relationships are among the most poorly understood, and ferns have long been thought to have few interactions with insects, although recent evidence suggests that these relationships are under-sampled and studied. Here, we present a near-chromosome genome assembly of the crambid moth *Neomusotima conspurcatalis,* a biological control agent employed in the management of the invasive vining fern *Lygodium microphyllum*. We use this novel genomic resource to explore the evolution of genome architecture across the Crambidae, revealing highly conserved genome structure across this family of moths. We also examine gene family evolution across the phylogeny and identify expansions in odorant receptor gene families that may be involved in the highly specific interaction of *N. conspurcatalis* with *L. microphyllum*. This work highlights the utility of genomics in biological control, and the utility of biological control in informing fundamental understanding of plant-insect interactions.

**Summary:** Plant-insect interactions form the basis of ecological communities and can be highly specific through long lasting co-evolution. Classical biological control provides excellent opportunities to investigate how insect and host genomes are shaped by co-evolutionary processes. We sequenced the genome of *Neomusotima conspurcatalis,* a biological control moth used to manage the invasive fern *Lygodium microphyllum.* We analyzed how the structure of the genome has evolved over time relative to other crambid moths and how expansions in gene families involved in odorant reception may be involved in co-evolution with its host. This work provides insight into herbivore interactions in seed-free vascular plants.

## INTRODUCTION

Plant-insect interactions are foundational ecological relationships within most terrestrial and freshwater aquatic ecosystems (Labandeira 2013; Yang and Gratton 2014). Co-evolution between host plants and associated insects (e.g., herbivores, pollinators) is expected to shape both genomes: for example, as plants evolve defenses to prevent herbivory, herbivores will evolve ways around these new defenses (Ehrlich and Raven 1964; Becerra et al. 2009; Futuyma and Agrawal 2009). Ferns are an ancient lineage of plants arising sometime in the Carboniferous or earlier (Testo and Sundue 2016; Pelosi et al. 2022) and are largely assumed to lack herbivore interactions (Brues 1920; Cooper-driver 1978; Hendrix 1980; Boch et al. 2016). Most notably, ferns have evolved strong biochemical defense mechanisms that appear to prevent herbivory, such as the constitutive expression of phytoecdysteroids, flavonoids, thiaminases, cyanogenic glycosides, and alkaloids (Castrejón-Varela et al. 2022). However, studies have since revealed a wide variety of fern-insect interactions (e.g., Balick et al. 1978; Hendrix and Marquis 1983; Santos et al. 2018; Suissa et al. 2024; Cariglino et al. 2025) and that these interactions have historically been under-reported (Mehltreter 2010).

The genomics and molecular mechanisms through which these interactions are established are still poorly understood, especially in non-model systems such as ferns. Studies on model plant-insect interactions have revealed the role of receptors and signaling cascades in herbivory (Howe and Jander 2008; Erb and Reymond 2019; Snoeck et al. 2022). For example, using a combination of genome-wide association studies (GWAS) and experiments examining gene expression at various stages of establishment, Nallu et al. (2018) characterized the dynamic evolution of the interaction between the plant *Arabidopsis thaliana* and butterfly *Pieris rapae.* Comparative genomic studies have used analyses of gene family evolution to pinpoint genes putatively involved in plant-insect interactions (e.g., Weng et al. 2025), particularly in loci involved in chemical recognition between insects and plant hosts (e.g., Kwak et al. 2023). Such results provide valuable insights into the genomics of ecological interactions, but are still rare/non-existent for fern-insect interactions.

Classical biological control is the use of co-evolved natural enemies (e.g., pathogens, parasites, predators) to reduce or control populations of invasive species (Briese 2000; Seastedt 2015). This management tool is perhaps the most applied concept within community ecology and is founded upon the Enemy Release Hypothesis - that a species is released from enemies upon introduction into a new range (Keane and Crawley 2002). The use of biological control to control invasive species provides an outstanding opportunity to investigate the molecular mechanisms and evolution of plant-insect interactions (Wilson 1965). Invasive plants have become ubiquitous members of global ecosystems (Pyšek et al. 2020) and are both economically (Diagne et al. 2021; Fantle-Lepczyk et al. 2022) and ecologically (Wilcove et al. 1998; Vilà et al. 2011) harmful. The establishment of biological control, however, remains a significant challenge, especially for plants that have evolved efficient defense responses (Pappas et al. 2017). Elucidating the molecular and genomic underpinnings of biological control establishment can lead to more effective management strategies and provide insight into the mechanisms shaping these highly specific relationships.

*Lygodium microphyllum* is a highly impactful invasive fern species introduced to Florida in the late nineteenth century, originally native to southeastern Asia, Australia, the Pacific Islands, and Africa. In its invaded range, *L. microphyllum* forms extensive rhizomatous growth, generating dense mats that smother and shade native ecosystems, disrupt natural fire regimes (Pemberton and Ferriter 1998; Smith et al. 2025), and cause canopy collapse (Foxcroft et al. 2017). This species forms dense populations in Florida and is classified as a United States Federal Noxious Weed (Smith et al. 2025) that poses threats to vulnerable natural ecosystems (Pemberton and Ferriter 1998; Foxcroft et al. 2013) and agricultural systems including pine logging activities and ranchlands (Florida Exotic Pest Plant Council Lygodium Task Force 2006; Koop 2009). *Lygodium microphyllum,* like many other ferns, is known to be rich in secondary metabolites (Garrison-Hanks 1998; Smith et al. 2016; Wheeler et al. 2021; Carrasco and Chambers 2025) which may inhibit or impede herbivory including biological control establishment.

Surveys for suitable biological control arthroprods for *L. microphyllum* identified several crambid moths as possible agents (e.g., *Austromusotima camptozonale, Neomustima conspurcatalis, Lygomusotima stria*; Goolsby et al. 2003). *Neomusotima conspurcatalis,* commonly known as the brown *Lygodium* moth or *Lygodium* defoliator moth, has proven to be a highly specific and a particularly promising avenue for managing invasive populations of *L. microphyllum* (Boughton and Pemberton 2009; Boughton and Pemberton 2012; Smith et al. 2014). Larvae of *N. conspurcatalis* feed on the leaflets of *Lygodium,* leading to foliar damage and eventual tissue death (Boughton and Pemberton 2009), with plants that have experienced herbivory showing limited regrowth. The moth has readily established field populations following initial introduction in 2008, with persistence throughout central Florida and spread into areas neighboring areas of initial release (Smith et al. 2014).

The crambid moths (Crambidae), commonly known as grass moths, are a large group of lepidopterans with over 10,000 species (Léger et al. 2021), including *N. conspurcatalis*. Crambid moths occupy a wide variety of habits and geography inhabiting all continents except for Antarctica. Many crambid moths are considered damaging agricultural pests (e.g., *Diatraea saccharalis,* Capinera 2023a; *Ostrinia nubilalis,* Capinera 2023b) but others are commonly employed as potential biological control agents due to their dietary specificity (e.g., *Niphograpta albiguttalis*, Tipping et al. 2014; *Acentria ephemerella,* Batra 1977). Crambid subfamilies specialize on a range of host plants across a phylogenetic breadth from bryophytes (e.g., Scopariinae) to angiosperms (e.g., Acentorpinae, Schoenobiinae, Crambiae; Fig. 1). Only two subfamilies in Crambidae, the Musotiminae and Hoploscopinae, are specialist herbivores on ferns and lycophytes (Léger et al. 2021). A growing wealth of genomic resources in this clade, including the recent publication of the genome of a member of the Musotiminae (*Musotima nitidalis*, Sterling et al. 2024), offer increasing opportunities to explore the evolutionary genomics of the interactions of crambids with their host plants.

**Figure 1.**
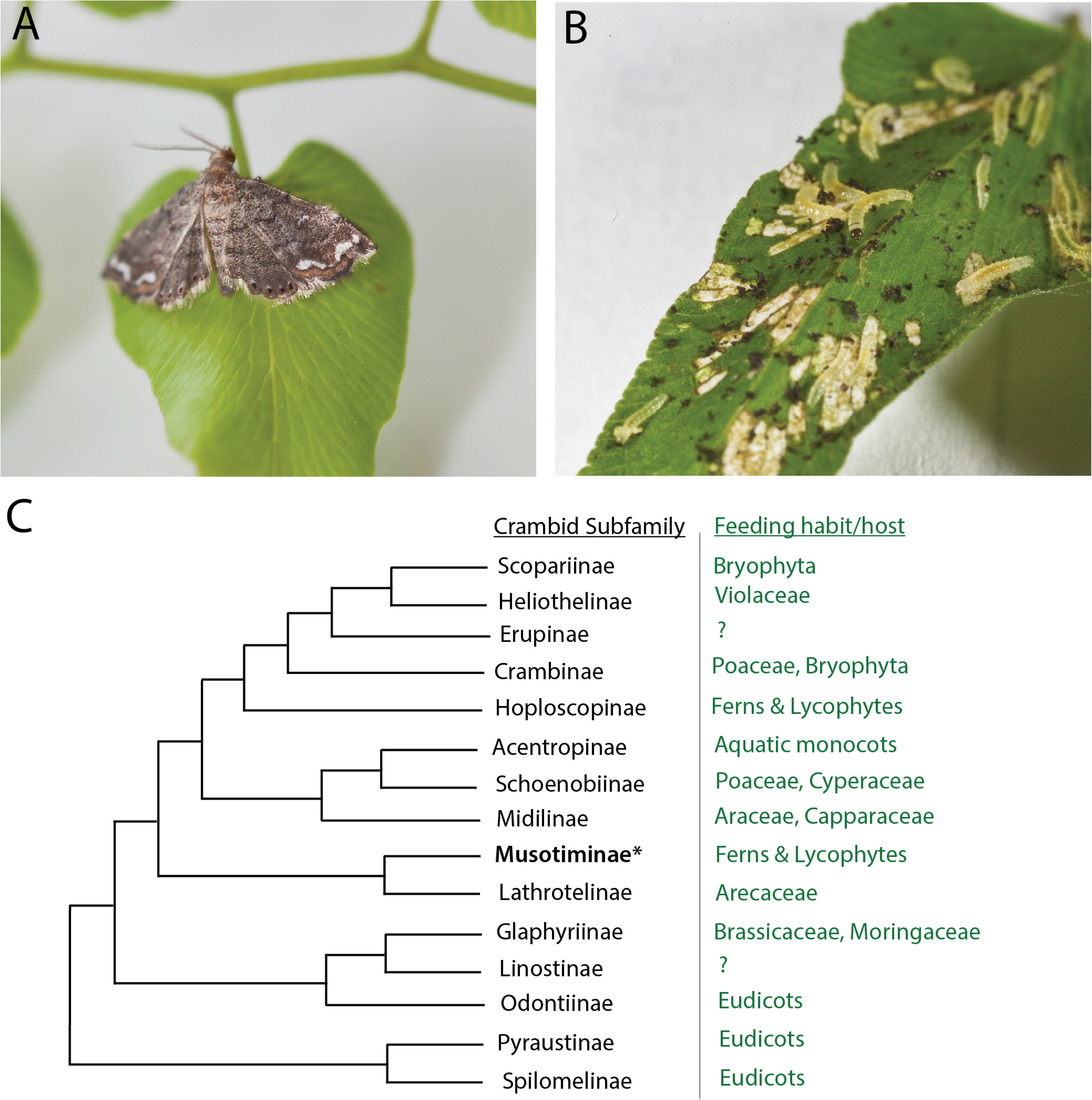
Photographs of *Neomusotima conspurcatalis* A) adult and B) larvae. C) Phylogeny of crambid subfamilies from (Léger et al. 2021) and corresponding host plant clade(s). Photographs taken by Melissa C. Smith.

Here, we sequenced the genome of *Neomusotima conspurcatalis*, producing a highly contiguous, near-chromosome level assembly. We combine this new genomic resource with a phylogenomic approach to investigate genome architecture and gene family evolution in the Crambidae, providing insight into the evolution of fern-insect interactions. Our analyses reveal highly stable genome architecture of crambid moths and inform how plant chemistry may shape the specificity of fern-insect interactions. Our genomic investigation of this biological control agent thus facilitates answering fundamental questions about the evolution of plant-insect interactions, while also informing on-going applied management strategies for invasive species such as *Lygodium microphyllum*.

## MATERIALS AND METHODS

### Sample Collection

Specimens were collected from the mass-rearing colony maintained at the United States Department of Agriculture (USDA) Invasive Plant Research Laboratory in Fort Lauderdale, Florida. Moths are reared for several generations from field-collected larvae and pupae (see Supplemental Table 1 for locations). For the collections, fourth-instar larvae and pupae were collected from the disease-free colony, placed into individual 2.5ml Eppendorf tubes, immediately submerged into liquid nitrogen, and stored at -80℃ until extraction (see Boughton and Pemberton 2012 for images of life stages). A voucher specimen from one of the field collection sites has been deposited in the University of Arizona Insect Collection (accession ID XXXXXXX).

### DNA Extraction

DNA extraction and sequencing was performed by the Arizona Genomics Institute (Tucson, AZ, USA). High molecular weight DNA was extracted from four pooled individual larvae which were homogenized in an extraction buffer with 0.1M Tris HCl buffer pH 8.0, 0.1M EDTA pH8, 1% SDS and Proteinase K at 50°C for 60 minutes. The mixture was centrifuged and aqueous phase transferred to a new tube to which 5M Potassium acetate was added, precipitated on ice, and centrifuged. After centrifugation, the supernatant was gently extracted with 24:1 chloroform:isoamyl alcohol. The upper phase was transferred to a new tube and DNA precipitated with 2/3 volume isopropanol. DNA was collected by centrifugation, washed with 70% ethanol, air dried, and dissolved thoroughly in 10 mM Tris-HCl followed by RNAse treatment. DNA purity was measured with Nanodrop, DNA concentration measured with the Qubit HS DNA assay (Invitrogen, MA, USA) and DNA fragment size validated by Femto Pulse System (Agilent, CA, USA).

### PacBio HiFi Sequencing

DNA was sheared to an appropriate size range (10-20 kb) using Megaruptor 3 (Diagenode, NY, USA) followed by SMRTbell cleanup beads. The sequencing library was constructed following the manufacturers’ protocols using the SMRTbell Prep kit 3.0. The final library was size-selected with AMPure PB beads. The recovered final library was quantified with a Qubit HS DNA assay (Invitrogen, MA, USA) and the size distribution checked on a Femto Pulse System (Agilent, CA, USA). The final library was cleaned with the Qiagen DNeasy powerClean Pro Cleanup (Qiagen, Netherlands), and then prepared for sequencing with the PacBio Revio SPRQ Sequencing kit for HiFi library, loaded on two Revio SMRT cells, and sequenced in CCS mode for 24 hours.

### Nuclear Genome Assembly & Annotation

We first mapped HiFi reads to the *Lygodium microphyllum* reference genome (Pelosi et al. 2025) with minimap2 v2.24-r1122 (Li 2018). Very few reads mapped to the *L. microphyllum* nuclear or chloroplast genomes (0.96% mapped to the nuclear genome, < 0.01% mapped to the plastome). We used the non-mapped reads as input for KMC v3.2.4 (Kokot et al. 2017) to generate a *k-*mer frequency spectrum with several values of *k* (*k*=17, 21, 31, 41, 51, 61), which we used as input for a custom R script (see Data Availability) to estimate genome size and heterozygosity.

We assembled the reads with hifiasm v0.16.1-r375 (Cheng et al. 2021) with the following modifications: we specified that there were 8 haplotypes present in our data (--n-hap 8), a haploid genome size of 500Mb (--hg-size 500m), and a *k*-mer size of 21 (-k 21). To ensure we did not assemble more than one representative haplotype, we checked the completeness of the assembly with compleasm v0.2.6 (Huang and Li 2023) with the lepidoptera_odb10 dataset. We ran purge_haplotigs v1.1.3 (Roach et al. 2018) and discarded contigs <1 Mb to remove any possible haplotypic duplications in the assembly. Telomeric repeats were identified with tidk v0.2.41 (Brown et al. 2025) with the lepidopteran telomeric repeat sequence (AACCT)*_n_*.

Amino acid sequences of predicted gene models (proteome data) for nine Crambidae and two Pyralidae moth genomes that represent a phylogenetic breadth of the family were downloaded from ENSEMBL, NCBI, or author-provided supplemental material in publications (Supplemental Table 2). These data were used as the input into the BRAKER3 v3.0.6 pipeline (Gabriel et al. 2024) to annotate the *Neomustomina conspurcatalis* nuclear genome assembly, which uses GeneMark-EP+ (Lomsadze et al. 2005; Brůna et al. 2020), DIAMOND (Buchfink et al. 2015), Spaln2 (Gotoh 2008; Iwata and Gotoh 2012), and AUGUSTUS (Stanke et al. 2006; Stanke et al. 2008). We assessed the completeness of the annotation with BUSCO v6.0.0 (Tegenfeldt et al. 2025) with the lepidoptera_odb10 dataset and OMArk v0.3.0 (Nevers et al. 2025) with the ancestral clade Obtectomera. MitoHiFi v2 (Uliano-Silva et al. 2023) was used to assemble the mitochondrial genome, which relies on MitoFinder (Allio et al. 2020) for annotation.

### Phylogenomics

We downloaded chromosome-scale genomes for species representing the phylogenetic breadth of the family and three outgroups. These included two Pyralidae (*Galleria mellonella, Amyelois transitella*), one Nymphalidae (*Heliconius sara*), and 14 Crambidae including two Acentropinae (*Elophila nymphaeata*, Boyes et al. 2024; *Parapoynx stratiotata,* Boyes et al. 2022), two Crambinae (*Chilo suppressalis*, Ma et al. 2020; *Diatraea saccharalis*), one Odontiinae (*Heortia vitessoides*, Law et al. 2022), two Pyraustinae (*Loxostege sticticalis*, Zhang et al. 2025; *Ostrinia nubilalis*, Ding et al. 2025), two Schoenobiinae (*Schoenobius gigantellus*, *Scirpophaga incertulas,* He et al. 2025), two Scopariinae (*Eudonia lacustrata,* Boyes et al. 2023; *Scoparia ambigualis,* Hutchinson et al. 2025), two Spilomelinae (*Maruca vitrata,* Wang et al. 2024; *Mecyna flavalis,* Sims et al. 2024), and one Musotiminae (*Musotima nitidalis,* Sterling et al. 2024; in addition to *Neomusotima conspurcatalis*) from ENSEMBL, NCBI, or author-provided supplemental material in publications (Supplemental Table 2). The genome annotation files (coding sequences, GFF) were also downloaded when available. Some genomes lacked or had erroneous annotations (*Scoparia ambigualis, Schoenobius gigantellus, Scirpophaga incertulas*); for these taxa we used the BRAKER3 pipeline as above to generate gene model predictions.

To generate a nuclear phylogeny and examine gene family evolution, we first clustered genes into putative gene families (i.e., orthogroups). Prior to clustering, we retained only the longest isoform for each gene using AGAT v1.5.1 (Dainat et al. 2025) or for genomes downloaded from ENSEMBL, only the primary isoform using a python script from Emms and Kelly (2015). OrthoFinder v3.1.0 (Emms and Kelly 2015; Emms and Kelly 2019; Emms et al. 2025) was used to cluster resulting peptide sequences into orthogroups. The corresponding coding sequences were extracted using SeqKit (Shen et al. 2016) for a total of 2582 single-copy orthogroups containing all taxa (100% taxon occupancy). For each orthogroup, we aligned the sequences with the codon-aware alignment tool MACSE v2.07 (Ranwez et al. 2018) and removed sites with less than 50% occupancy with trimAl v1.5.rev0 (Capella-Gutiérrez et al. 2009). Maximum-likelihood gene trees were constructed using IQTREE3 v3.0.1 (Wong et al. 2025) with 1000 ultrafast bootstraps (Hoang et al. 2018) after estimating the best-fitting substitution model with ModelFinderPlus (Kalyaanamoorthy et al. 2017) for each gene. These gene trees were input to ASTRAL IV (Zhang et al. 2025) to generate a species tree using the multi-species coalescent model.

### Gene Family Evolution

To examine how gene families evolved along the phylogeny and whether changes in gene family size were associated with herbivory strategy, we used CAFÉ v5.1 (Mendes et al. 2021). First, we generated a species tree with branch lengths proportional to the number of substitutions per site by concatenating the 2582 single-copy orthogroups into a single phylogenetic matrix which we used as input to IQTREE3 v3.0.1 (Wong et al. 2025) where we used the ASTRAL species tree as a topological constraint and estimated the best-fitting substitution models for each locus with ModelFinderPlus (Kalyaanamoorthy et al. 2017). We then used treePL v1.0 (Smith and O’Meara 2012) to estimate divergence times with two calibration points: a fossil calibration at stem Pyrallidae (47.8-56.0 MY) and a secondary calibration for the divergence between butterflies and moths (95.9-119.4 MY; Kawahara et al. 2019). We performed three cross-validation runs for each tree and determined the optimal smoothing parameter based on the lowest chi-squared value. This ultrametric tree was used as the backbone phylogeny in CAFÉ v5.1 (Mendes et al. 2021) to estimate significant expansions and contractions and rate shifts in gene family evolution. For CAFÉ, we estimated a single λ across the tree with one rate category (Base, one γ), two rate categories (k=2), and three rate categories (k=3). For all three parameter sets, we specified a poisson distribution for the root frequency distribution, and for the Base model we first estimated an error model to account for mis-assemblies and mis-annotations prior to running the analysis (empirical error model estimation is not available for multiple gamma values). Three independent runs of each parameter set were inspected to verify convergence. The best-fitting model was determined using a likelihood ratio test. We used CafePlotter (https://github.com/moshi4/CafePlotter) to parse and visualize the results of CAFÉ. To assign putative functions to gene families, we used BLASTP v2.13.0 (Camacho et al. 2009) to identify sequence homology between the predicted *Neomusotima conspurcatalis* proteins and *Drosophila melanogaster* (R6.65) downloaded from FlyBase (release FB2025_04; Öztürk-Çolak et al. 2024). The BLAST hit with the lowest E-value was retained for each *N. conspurcatalis* gene, and the corresponding *D. melanogaster* gene information was carried over as annotation to the orthogroup containing the *N. conspurcatalis* gene.

### Genome Architecture

To examine the evolution of macrosyntenic patterns of genome architecture across Crambidae, we used GENESPACE v1.3.1(Lovell et al. 2022). The same 18 genomes used for the nuclear-sequence based phylogenomics were input to this pipeline, which relies on OrthoFinder v2.5.5 (Emms and Kelly 2015; Emms and Kelly 2019) and MCScanX v1.0.0 (Wang et al. 2012). We also used D-GENIES (Cabanettes and Klopp 2018) to examine pairwise comparisons of *Musotima nitidalis* and *Neomusotima conspurcatalis.* To further interrogate microsynteny, we used the R package syntenet v1.6.1 (Almeida-Silva et al. 2023). Using the same 18 genomes as for GENESPACE, we used DIAMOND v2.1.13.167 (Buchfink et al. 2015) to perform all-by-all sequence similarity searches for the proteomes and filtered the results by only retaining hits with >70 percent identity scores and longer than 30 amino acid residues as in MacGuigan et al. (2025). We then inferred syntenic networks within syntenet, which implements the MCScanX algorithm to detect synteny. The resulting syntenic network was then clustered with default parameters and subjected to phylogenomic profiling following Almeida-Silva et al. (2023). We further used the function *‘binarize_and_transpose()’* to generate a binary matrix of the profiles, with which we ran IQTREE3 v3.0.1 (Wong et al. 2025) with the MK+FO+R model and 1000 ultrafast bootstraps (Hoang et al. 2018).

## RESULTS

### Sequence Data

A total of 8,700,936 HiFi reads with a read N50 of 7.2Kb equalling 71.78Gb of data were generated. Assuming a genome size around 500Mb (based on other genomes in the family), we expect approximately 140x haploid genome coverage. When we examined the *k*-mer spectra, the homozygous peak was ∼150x *k*-mer coverage, but there was also a large heterozygous peak around 18x coverage, which is consistent with the detection of 8 individual haplotypes within the pooled sample (Supplemental Figure 1). Despite the large heterozygous peak, we estimated that the haploid genome of *Neomusotima conspurcatalis* was between 345-412Mb, which is similar to the haploid genome size of *Musotima nitidalis* (555Mb), the only other assembled genome for this subfamily.

### Genome Assembly and Annotation

Using the --n-hap flag (which was designed for polyploid genome assembly), we were able to recover a single representative haplotype that was highly contiguous and had low duplication rates. The initial assembly from hifiasm prior to purging duplications was 465.6Mb and contained 390 contigs, with a contig N50 of 16Mb and 99.74% complete BUSCOs (3.5% duplicated). After purging duplicate contigs and removing short sequences (<1Mb), we attained a final assembly for *Neomusotima conspurcatalis* that was 441.1Mb in length contained in 40 contigs that had a contig N50 of 16Mb and 99.77% complete BUSCOs (0.81% duplicated). We identified 18 putative telomere-to-telomere contigs that may correspond to full chromosomes, and 17 contigs had high frequencies of telomeric repeats on one end (Supplemental Figure 2).

The nuclear genome annotation contained a total of 18,663 protein-coding genes and 21,110 transcripts and was highly complete based on BUSCO (98.3% complete genes, 96.5% single-copy, 1.8% duplicated BUSCOs; Table 1) and OMark scores (96.98% complete hierarchical orthologous groups [HOGs], 86.55% single, 10.53% duplicated). The phylogenetic placement of HOGs was largely consistent (77.15%, n=14399) with only 5.19% showing inconsistent placement. There were an average of 5.5 exons per coding sequence, and most genes were multi-exonic with only 4365 genes containing a single exon (23.3%).

**Table 1.** Genome assembly and annotation statistics for *Neomusotima conspurcatalis* v1.

| Data Type | Genome Metric | Value |
| --- | --- | --- |
| Assembly | Total Assembly Size (bp) | 441,162,612 |
|  | # of Contigs | 40 |
|  | Contig N50 | 16,002,885 |
|  | Longest Contig | 24,778,440 |
|  | # Putative T2T Chromosomes | 18 |
|  | Genome BUSCO (Lepidoptera_odb10, n=5286) | 99.77% complete (98.92% single, 0.81% duplicated) |
| Annotation | % Repetitive Elements | 44.58% |
|  | # Protein-coding genes | 18,663 |
|  | Proteome BUSCO (Lepidoptera_odb10, n=5286) | 98.3% complete (96.5% single, 1.8% duplicated) |
|  | Proteome OMARK (Obtectomera, n=6779) | 96.98% complete (86.44% single, 10.53% duplicated) |

### Phylogenomics

The final nuclear matrix contained 2,582 genes totalling 4,607,347bp with an average gene length of 1,784bp (range: 252-16,184). The resulting ASTRAL topology had strong support at every node (local posterior probably = 1), but there was substantial variation in gene-tree species-tree discordance (Figure 2A). The support for an alternative topology ranged from 0.13% to 30.26% of gene trees (Figure 2A). The ASTRAL topology recovered all subfamilies as monophyletic, with Musotiminiae sister to a clade composed of Schoenobiinae and Acentropinae. Together, this clade was sister to Scopariinae + Crambinae. Successively sister to this larger clade were *Heortia vitessoides* (Odontiinae), Pyraustinae + Spilomelinae, Pyralidae, and *Heliconius sara* (Nymphalidae). Of particular interest to our work, there was particularly high conflict at the node subtending the Musotiminae (percent of gene trees supporting alternate topologies 1 = 28.19% and 2= 30.26%); the branch leading to this node was especially short.

**Figure 2.**
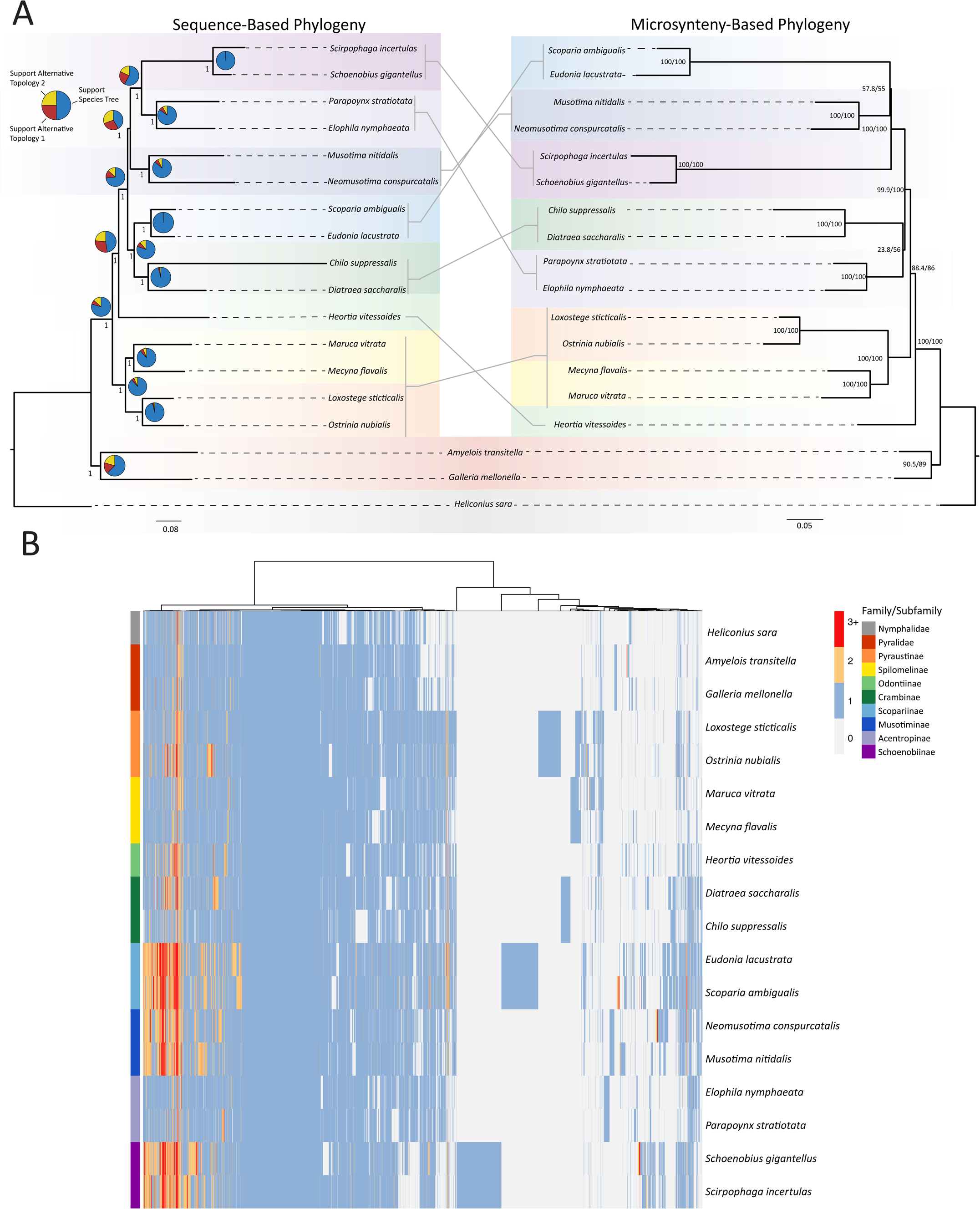
Phylogenomics of Crambidae reveals discordance. A) ASTRAL sequence-based phylogeny (left) generated from 2582 single-copy orthologs with local posterior probability node support values. Pie charts show the proportion of genes that support the species-tree topology (blue) and alternative topologies (yellow and red). Branch lengths are proportional to the number of substitutions per site. Microsynteny phylogeny generated from 16238 binary microsyntenic cluster profiles with SH-aLRT/ultrafast bootstrap node support values. Lines between phylogenies connect taxa, showing the relative placement in each tree. B) Heatmap of syntenic clusters, with taxa ordered based on ASTRAL sequence-based phylogeny in (A). Rows represent species and columns represent microsyntenic clusters, colored on the number of genes in each cluster. Gray space indicates the absence of genes in that cluster for that species. Dendrogram depicts the clustering based on Ward’s minimum variance hierarchical clustering.

The microsyntenic phylogeny was generated from 16,238 binary microsyntenic cluster profiles and nodes had varying levels of support as measured by SH-aLRT and ultrafast bootstrap values (Figure 2A). The topology was substantially different from the sequence-based phylogeny: the phylogeny recovered Musotiminae sister to Scopariinae, but with low support (SH-aLRT = 57.8, ultrafast bootstrap = 55). However, a clade composed of Musotiminae, Schoenobiinae, and Scopariinae was strongly supported (SH-aLRT = 100, ultrafast bootstrap=100). Successively sister to that clade were Crambinae + Acentropinae, Pyraustinae + Spilomelinae, *Heortia vitessoides* (Odontiinae), Pyralidae, and *Heliconius sara* (Nymphalidae) (Figure 2A). Notably, the genomes in the clade composed of Musotiminae, Scopariinae, and Schoenobinnae in this analysis were all annotated with BRAKER in some way: we used BRAKER to annotate *Neomusotima conspurcatalis, Scoparia ambigualis, Schoenobius gigantellus,* and *Scirpophaga incertulas,* and the annotations downloaded from ENSEMBL for *Musotima nitidalis* and *Eudonia lacustrata* are labeled with ‘BRAKER’, although the exact annotation method used is unclear for these latter two taxa.

### Genome Architecture

We recovered highly conserved macrosynteny across the crambid moth genomes analyzed (Figure 3). In most cases, entire chromosomes were conserved as syntenic blocks (i.e., the gene order along entire chromosomes were conserved across taxa, Figure 3). There are several instances of structural variants such as inversions, translocations, and, especially in Schoenobiinae, putative chromosomal fusions (Figure 3). The genomes of *Diatraea saccharalis* and *Chilo suppressalis* (Crambinae) have substantial differences in synteny compared to the other Crambidae, with several translocations in both taxa. There are several subfamily-specific syntenic clusters that are mostly in single copy (e.g., Schoenobinnae; Figure 2B). Of the microsyntenic clusters shared among the genomes analyzed, there was variation in copy number (Figure 2B). There are several clusters that are mulit-copy in Scopariinae, Musotiminae, and Schoenobiinae (3+ copies); this pattern appears to be mostly specific to these taxa, with most syntenic clusters in single copy across the Crambidae (Figure 2B).

**Figure 3.**
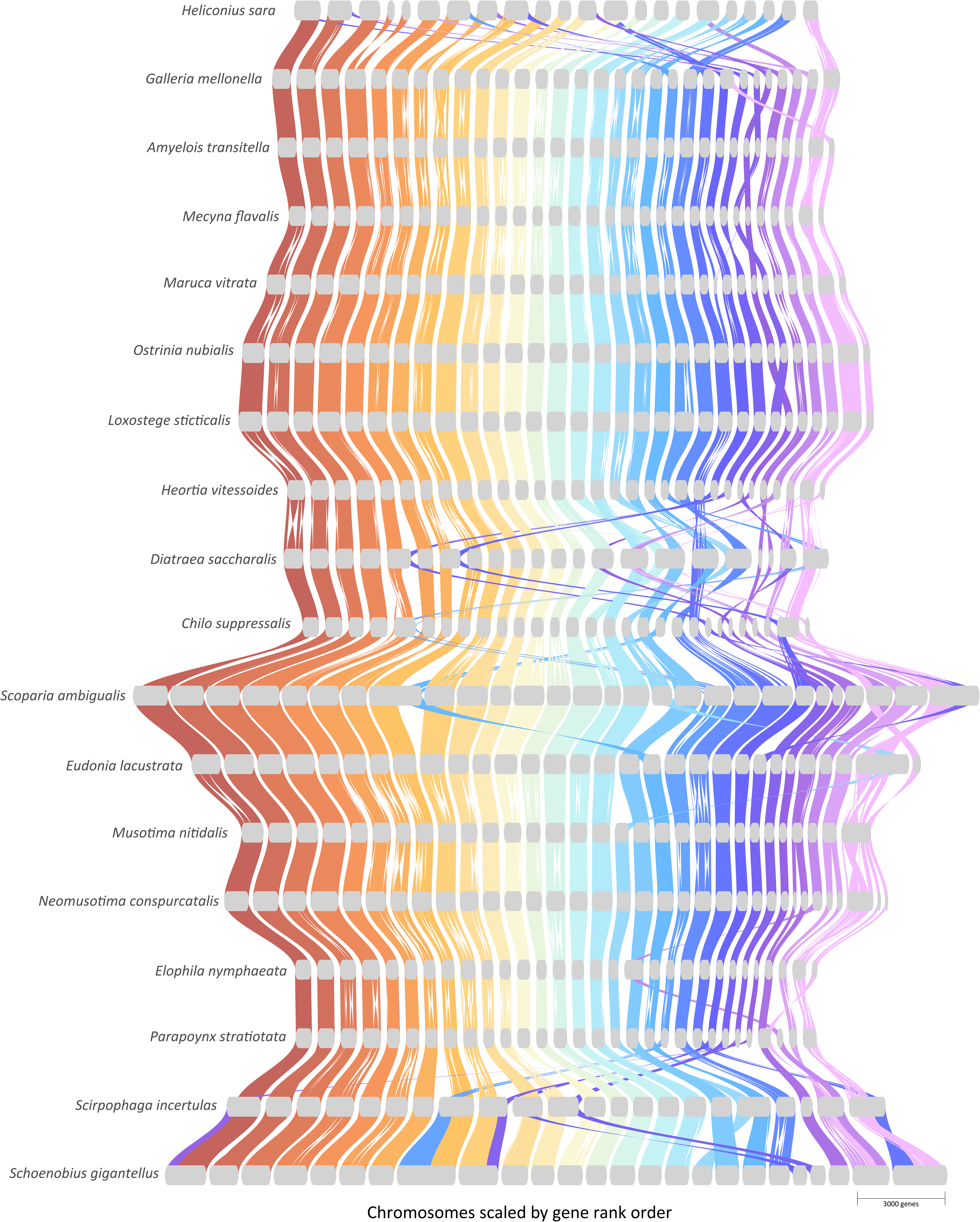
Macrosyteny of the Crambidae is highly conserved. Riparian plot of the Crambidae, where colored ribbons depict blocks of conserved synteny between species. Chromosomes are ordered based on *Musotima nitidalis* and scaled by gene rank order. The scale bar represents 3000 genes.

Comparing just the genomes of *Musotima nitidalis* and *Neomusotima conspurcatalis*, macrosynteny was largely conserved. Despite the clearly conserved 1:1 synteny between the two genera, there were several inversions (e.g., two inversions on each end of *Musotima* chromosome 6 and *Neomusotima* contig ptg000054l) (Supplemental Figure 3). Nearly all of the contigs we assembled in the *N. conspurcatalis* genome had syntenic hits to a single *Musotima* chromosome with some exceptions: ptg000006l and ptg000047l to *Musotima* chromosome 27, ptg00108l and ptg000136l to *Mustoima* chromosome 29, and ptg00007l, ptg000033l, and ptg000071l to the *Musotima* Z chromosome (Supplemental Figure 3).

Perhaps the most dynamic evolution of gene order was observed in the Z chromosome (Supplemental Figure 4). Genes on the *Musotima nitidalis* Z chromosome generally had orthologs on two chromosomes across the crambids. For example, the *M. nitidalis* Z chromosome has orthologs to the Z chromosome and autosome 29 from *Eudonia.* Across all chromosomes, the Z chromosome exhibited the greatest number of rearrangements such as inversions. For the *Neomusotima conspurcatalis* genome, we recovered three contigs with syntenic relationships to the *M. nitidalis* Z chromosome.

### Gene Family Evolution

Across the 18 genomes, a total of 297,759 genes were clustered into 16,290 orthogroups, with only 4.6% of genes remaining unassigned. Of these orthogroups, 1.3% were species-specific. For *N. conspurcatalis,* 92.7% of predicted genes were clustered into orthogroups, with 0.8% of genes assigned to a species-specific orthogroup. The best-fitting model of gene family evolution was with two rate categories (k=2, likelihood ratio test, *P* < 0.005). A total of 11392 gene families were retained after filtering for families that were not present at the root of the phylogeny. Of these, 314 gene families had significant changes in size (i.e., underwent an expansion or contraction) somewhere on the phylogeny (Figure 4). Along the branch subtending the Musotiminae, there were 27 significant expansions and 8 significant contractions. The expanded gene families had putative orthologs involved in DNA binding, protein degradation and chitin development Along the terminal branch leading to *Neomuostima conspurcatalis* there were 62 significant expansions and 33 significant contractions; these include gene families with sequence homology to genes involved in DNA binding, metabolism, and receptors. Two gene families of particular interest that underwent a significant expansion in this latter category were odorant chemoreceptors that contained putative homologs of the *Drosophila melanogaster* genes *Or85b* and *Or85c*, which code for odorant chemoreceptor that are sensitive to volatile organic compounds (Figure 4B).

**Figure 4.**
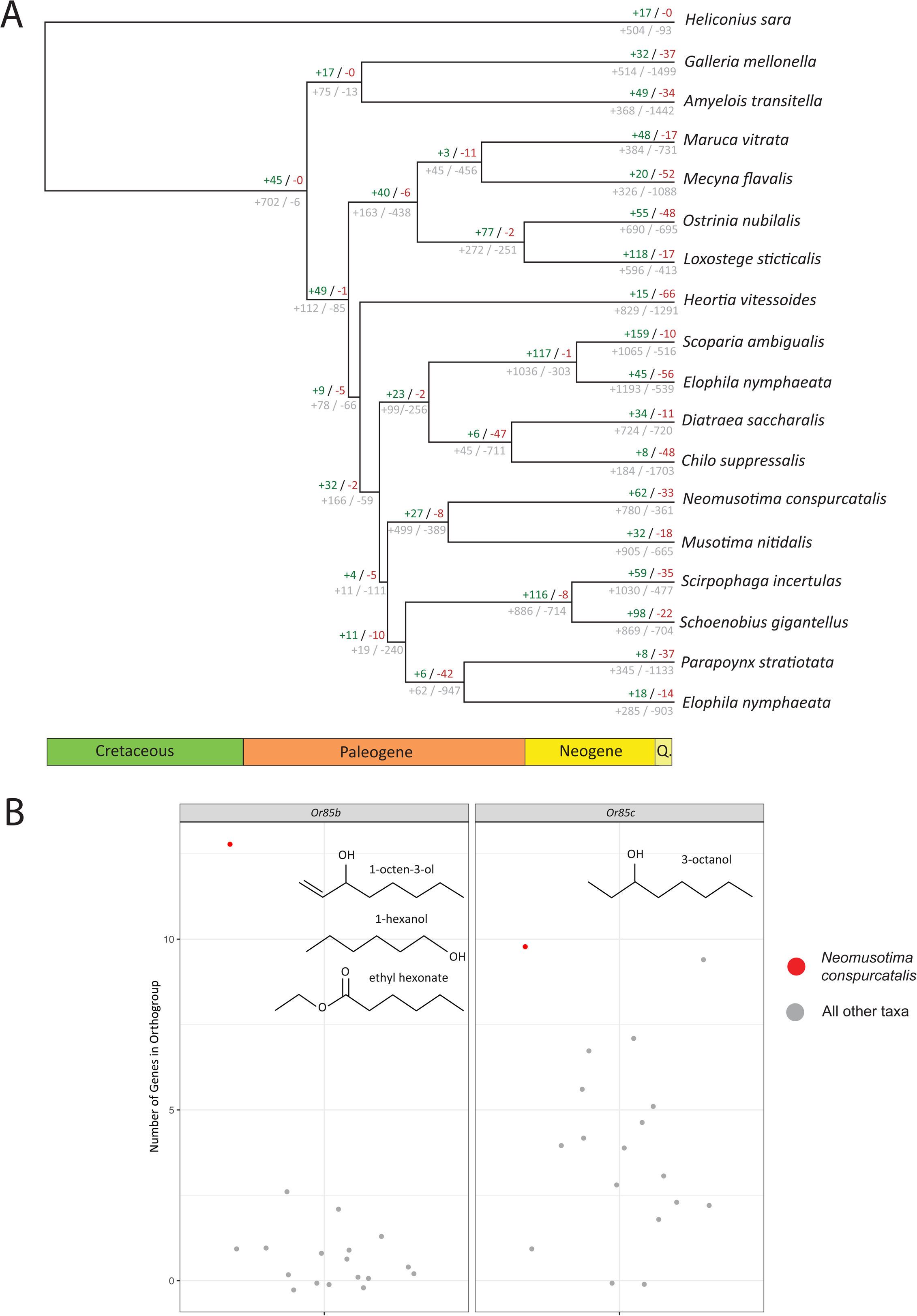
Gene family evolution in the Crambidae. A) Branches are annotated with the overall number of expansions / contractions (gray) below and significant expansions (green) / contractions (red) above. B) Two gene families with significant sequence homology to *Drosophila melanogaster* Or85b (left) and Or85c (right). Number of members in each moth genome (gray) and *Neomusotima conspurcatalis* (red). Chemical structures of the volatile organic compounds (VOCs) that each odorant receptor detects are shown in each plot.

## DISCUSSION

The genomics of plant-insect interactions are particularly understudied in non-model seed-free plants, such as ferns. The Crambidae are one of the largest clades of moths with over 10,000 species and include taxa with unusual feeding specificity on ferns and lycophytes (Léger et al. 2021). Their impacts on agriculture and use as biological control agents, together with their increasing genomic resources, makes the Crambidae an excellent system for investigating the evolutionary genomics of plant-insect interactions. To this end, we assembled a near-chromosome scale genome for the biological control agent *Neomusotima conspurcatalis*, which feeds nearly exclusively on the invasive fern *Lygodium microphyllum* (Boughton et al. 2009). We used this novel resource to explore the evolution of genome architecture of the crambids and investigate the genomic underpinnings of their specialization on ferns.

*Genome-Scale Phylogenetics of the* Crambidae The phylogenetic relationships of Crambidae have been unclear despite the efforts of several researchers (Roesler 1973; Kuznetzov and Stekolnikov, 1979; Yoshiyasu 1985; Solis and Maes 2002; Solis 2007), although recent molecular analyses have begun to shed light on these relationships (Regier et al. 2012; Léger et al. 2021). Both Regier et al. (2012) and Léger et al. (2021) recovered Musotiminae sister to the rest of the “CAMMSS clade” (Crambinae, Acentropinae, Midilinae, Musotiminae, Schoenobinnae, and Scopariinae), although Léger et al. (2021) also included samples from Lathrotelinae, which formed a clade with Mustominae and together were sister to the rest of the CAMMSS clade. Interestingly, Léger et al. (2021) also found that Hoploscopinae, the only other subfamily to specialize to pteridophytes in Crambidae, is nested within the CAMMSS clade, suggesting a convergent strategy of feeding on ferns.

Here we used both traditional phylogenomic methods (sequence-based) and newly developed microsyteny-based approaches to the crambid phylogeny, and recovered vastly different topologies. Given that coding sequences typically make up a small proportion of the genome, approaches that exploit more of the genome to infer phylogenetic relationships may greatly improve our understanding of evolution. For example, genome architecture, such as the physical position of genes relative to each other, may be a powerful character to use in phylogenetic reconstruction (Zhao and Schranz 2019; Zhao et al. 2021; Steenwyk and King 2024). From the sequence-based analyses, we found that Musotiminae was nested within the CAMMSS clade (rather than sister to the rest), was sister to a clade composed of Schoenobinnae + Acentropinae, and Scopariinae and Crambinae were together sister to the rest of the CAMMSS clade (Figure 2). Using a microsynteny approach, we recovered an altogether different topology, with Musotiminae sister to Scopariinae, Schoenobinnae sister to that clade, and a clade composed of Acentropinae + Crambinae sister to the rest of the CAMMSS clade. While both topologies conflict with (Regier et al. 2012; Léger et al. 2021), the placement of Musotiminae lacks strong support: in the sequence-based phylogeny, there is a high level of gene-tree species-tree conflict at the node subtending Musotiminae (Figure 2; although LPP = 1) and the same node has low support (SH-aLRT = 57.8, ultrafast bootstrap = 55) in the microsynteny analysis. Interestingly, the topology recovered using the microsynteny analysis is most similar to Yoshiyasu (1985), with Musotiminae sister to Scopariinae.

Importantly, however, the microsynteny approach may be highly sensitive to annotation quality and differences in annotation pipelines (Steenwyk and King 2024). It is possible that the highly supported clade we recovered of Musotiminae, Scopariinae, and Schoenobinnae in our microsynteny analysis is an artifact of different annotation methods: we annotated *Neomusotima conspurcatalis, Scoparia ambigualis, Schoenobius gigantellus,* and *Scirpophaga incertulas* and the annotations downloaded from ENSEMBL for *Musotima nitidalis* and *Eudonia lacustrata* are labeled with ‘BRAKER’ (the precise annotation pipeline for these taxa is unclear). Further testing will be needed to determine if the relationships we recovered in the microsynteny analysis are a true biological feature of these genomes, or arise from a bias in the annotations. Additional genomic sampling of other Crambidae subfamilies will enable further clarification of these relationships. For example, Léger et al. (2021) found Lathrotelinae sister to Musotiminae, but a chromosome-scale genome for this clade was not available and therefore could not be included in our analyses. Further sequencing of other subfamilies will undoubtedly provide insight into the evolutionary history of this clade.

The timing of the divergence among taxa may further provide insight into co-evolutionary dynamics between the insect and plant. We recovered a divergence time between *Neomusotima* and *Musotima* of around 34.45 MYA. While *Musotima nitidalis* is a generalist herbivore feeding on Pteridaceae, Dennstaedtiaceae, Dryopteridaceae, Blechnaceae, and Asplenaiceae (Sterling et al. 2024), *Neomusotima* is nearly exclusive to *Lygodium microphyllum*. The divergence between Lygodiaceae and the rest of the ferns occurred ca. 174 MYA (Pelosi et al. 2022), with the earliest *Lygodium* species originating in the Late Cretaceous (*Lygodium cretaceum,* Debey and Ettingshausen 1859). A species closely resembling *L. microphyllum* and its sister species *L. reticulatum* is known from the fossil record in Australia from the Oligocene - early Miocene (Rozefelds et al. 2017). The overlap in the divergence times of *Neomusotima* and *Musotima* (ca. 39 MY) and *L. microphyllum* and *L. reticulatum* from other *Lygodium* species (∼51 MY; Testo and Sundue, 2016) may support a long-term co-evolutionary relationship between the two species. Interestingly *Neomusotima* feeding does not differ between genotypes of *L. microphyllum* that are genetically divergent (Goolsby et al. 2003; Boughton and Pemberton 2009; Boughton and Pemberton 2012; Smith et al. 2014; Pelosi et al. 2024). A robust phylogenetic framework of the Musotiminae will provide valuable insight into the co-evolution of these moths with their host ferns. More specifically, given that functional traits may (but not always) be phylogenetically conserved, phylogenies can be used to identify new specific biological control agents.

### Evolution of Genome Architecture in the Crambidae

As the assembly of chromosome-level genomes becomes accessible for a greater number of non-model species, we can now study the evolution of the physical structure of the genome over evolutionary time. In arthropods, synteny is largely conserved (Engström et al. 2007; Tolman et al. 2023), especially in the Lepidoptera (Pringle et al. 2007; Traut et al. 2023). Our results are largely congruent with the existing literature: genome structure of this clade of moths is also well-conserved. In several species in our analysis spanning > 90 MY of evolution, there are syntenic blocks that span the entirety of chromosomes, meaning that the gene order along the chromosome is conserved across species. Most other studies examining crambid moth synteny have used pairwise methods to assess syntenic relationships (Ma et al. 2020; Law et al. 2022; Ding et al. 2025, but see Zhang et al. 2025 for a four-way comparison); by examining syntenic relationships using 18 highly-contiguous genomes, we find patterns of synteny that were not previously not apparent and can trace syntenic blocks across evolutionary time. For example, our analysis revealed that the evolution of syntenic blocks on the Z sex chromosome is highly dynamic compared to the mostly conserved synteny of the autosomes. There are instances where Z chromosome blocks are translocated to autosomes (e.g., in *Diatrea, Galleria, Paraponyx*) and putative fission in *Scirpophaga incertulas* and *Schoenobius gigantellus*. In contrast with most of the other crambid genomes, *Diatraea saccharalis* and *Chilo suppressalis* (both in the subfamily Crambinae) showed the most dynamic genome evolution with multiple translocations and rearrangements. *Schoenobius gigantellus* and *Scirpophaga incertulas* (Schoenobiinae) have substantially larger genomes compared to the other crambids (haploid size ca. 700Mb-1Gb versus ca. 500Mb); the change in size does not appear to be driven by genome duplication (polyploidy), as there is a lack of duplicated syntenic regions and BUSCOs. Interestingly, these two genomes also show evidence for chromosomal fusions, which are largely lacking in the other crambid moth genomes.

At a finer scale, a pairwise comparison between *Neomusotima conspurcatalis* and *Musotima nitidalis* reveals several structural variants between the two reference genomes. For example, there is a large inversion between *Musotima* chromosome 20 and *Neomusotima* contig ptg000056l and two inversions on each end of *Musotima* chromosome 6 and *Neomusotima* contig ptg000054l (Supplemental Figure 3B, C). Further study will be required to clarify whether these variants are fixed in *Neomusotima conspurcatalis* relative to *Musotima nitidalis.* With this newly sequenced genome, it is now possible to ask questions such as: what is the evolutionary relevance of inter- and intraspecific variation in chromosomal rearrangements? Do these variants play roles in reproductive isolation and/or local adaptation?

### Insights into Fern-Insect Interactions From Gene Family Evolution

Biotic interactions between insects and their host plants shape the evolution of both genomes (Gloss et al. 2013; Gloss et al. 2019). Plant chemistry is largely considered to be a defining factor in other stages of plant-insect interactions (e.g., Ehrlich and Raven 1964; reviewed by Nishida 2014; Dyer et al. 2018). For example, the differentiation of the host and non-host plants and oviposition is widely recognized as occurring through chemical cues such as the recognition of volatiles through odorant receptors (Renwick and Chew 1994; Bengtsson et al. 2006; Clavijo McCormick et al. 2014). Interrogating gene family evolution can be a powerful method to identify candidate genes involved in key evolutionary transitions, such as host specificity. For example, studies in *Drosophila* have revealed that chemosensory gene families are likely key players in the specificity of recognition: odorant-binding protein (*OBP*), odorant receptor (*Or*), and gustatory receptor (*Gr*) gene families have undergone rapid evolutionary change (McBride 2007; Vieira et al. 2007). In the Lepidoptera, the number of *Or* and *Gr* genes has greatly increased since the origin of the clade, although their DNA sequences are largely under strong purifying selection (Engsontia et al. 2014). *Or* genes are known to play particularly important roles in host specialization. For example, Liu et al. (2020) found that the specificity of the moth *Plutella xylostella* on crucifers was only established through the detection of isothiocyanates produced by the host plant through the odorant receptors *Or35* and *Or49*. Nearly nothing is known about how these interactions are established or the role of *Or* genes in seed-free vascular plants (i.e., pteridophytes) interactions with their insect herbivores.

Our observations of gene family expansions in *N. conspurcatalis* appear to align with a recent work by Smith et al. (2016) and Wheeler et al. (2021) on the chemical landscape of fern-insect interactions in *Lygodium.* By comparing the make-up of volatile organic compounds (VOCs), which are known to play a role in recruiting specialist herbivores (Renwick and Chew 1994; Bengtsson et al. 2006; Clavijo McCormick et al. 2014), among congeners in *Lygodium*, and integrating this with oviposition behavior of *N. conspurcatalis,* they found that *N. conspurcatalis* oviposition was highest on plants with similar volatile profiles to *L. microphyllum* regardless of phylogenetic distance. Three VOCs made up more than 75% of the volatile profile of *L. microphyllum*: 1-octen-3-ol (32.4%), 3-octanone (25.7%), and sativene (20.3%). While 1-octen-3-ol was detected in the six *Lygodium* species analyzed, the proportion of this compound in *L. microphyllum* was significantly different from *L. oligostachyum* and *L. palmatum* (Wheeler et al. 2021). Interestingly, we identified significant expansions in two *N. conspurcatalis* gene families with sequence homology to *Drosophila melanogaster Or85b* and *Or85c* genes known to respond to VOCs along the terminal branch leading to *N. conspurcatalis* (Figure 4). *Or85b* is involved in odorant responses of 1-octen-3-ol, 1-hexanol, and ethyl hexonate (Hallem and Carlson 2006), while *Or85c* is involved in responses to 1-octanol (Vosshall et al. 2000). This expansion may have been involved in the evolution of specificity between *N. conspurcatalis* and *L. microphyllum*, although further functional assays will be required to test for this relationship. For example, we can use an integrated transcriptomic and metabolomic approach to determine when these genes are expressed in *N. conspurcatalis* and in response to what types of compounds produced by *L. microphyllum.* Additional sequencing in other moths (such as those in Heliospinae) with specificity on ferns provide opportunities to examine whether there is convergence on similar molecular pathways. The *N. conspurcatalis* genome generated here provides a new resource for understanding the evolutionary genomics of ecological interactions.

### Application and Utility of Genomics in Biological Control

Genomics is increasingly being recognized as a valuable tool in the biological control community (Leung et al. 2020). As the accessibility of genome sequencing continues to expand, biological control genomics has the potential to improve identification, prioritization, post-release impact, and potential use in future genetic biological control applications. For example, field surveys of establishment can be improved through the use of molecular-based identification of whole samples when taxa are difficult to identify (e.g., cryptic species) or using environmental DNA (eDNA; Kirtane et al. 2022; Bell et al. 2024), which may better monitor biological control populations without requiring/needing for the visible presence of the biological control. Reference genomes in particular facilitate functional genomic approaches (e.g., quantifying gene expression, QTL mapping, GWAS) to identify loci associated with traits (e.g., pest-suppression, climate adaptation, fecundity and mass rearing, Leung et al. 2020). Knowing the genetic basis of these traits can inform how trait values are impacted by introduction pathways and bottlenecks, and which traits might be most advantageous in a particular environment for which to select. For example, quantitative traits controlled by few loci of large effect may be subject to more fluctuation in trait value during the colonization process than traits controlled by many loci of small effect (Dlugosch et al. 2015). Genomics allows researchers to identify how traits are governed by the genome, and provide insight into how trait values will shift with artificial selection. Furthermore, debate over the use of genetic modification (e.g., RNAi, CRISPR-Cas9) to improve biological control efficacy has highlighted the utility of detailing genome-phenotype relationships (Webber et al. 2015). For the *Neomusotima* genome presented here, an important future use includes identifying loci involved in and selecting for increased cold and freezing resistance. *Lygodium microphyllum* is expected to continue shifting northward (Pelosi et al. 2025) and the source population of the *Neomusotima* colony maintained by the USDA is from a largely tropical region in the center of its distribution (Australia). What regions of the genome are associated with cold and freezing tolerance? What is the level of standing genomic variation in these regions? Could artificial selection for increased tolerance provide a way for *Neomusotima* to track its host northward?

Evolutionary genomics, broadly, has the potential for immense predictive power (e.g., Long et al. 2025; Ortiz-Barrientos et al. 2025); harnessing and integrating techniques such as landscape genomics and genotype-environment association analyses for biological control can aid in identifying from where in the native range samples should be sourced and if loci under selection from environmental variables are shared or differ. For example, Barker et al. (in prep), found that loci associated with climate adaptation in the biological control weevil *Eustenopus villosus* largely differed between the native and introduced range. Application of spatial landscape genomics and genotype-environment associations have further been used to predict how genetic variation will vary over space under future climate scenarios (Fitzpatrick and Keller 2015). A similar approach may be employed to identify the (mis-)match of native range genotypes to the environmental conditions in the putative introduced range. A landscape genomic approach to *Neomusotima* could shed light on which region(s) in the native range to source new populations such that adaptive genomic variation is best-fit for the climate in the invaded range.

## AUTHOR CONTRIBUTIONS

J.A.P.: conceptualization, data analysis, writing + revising T.R.C.: data analysis

A.R.P.: data analysis

M.C.S.: conceptualization, sample collection, writing + revising K.M.D.: conceptualization, writing + revising

## ACKNOWLEDGEMENTS

We thank Trevor Krabbenhoft for valuable discussion on resolving haplotypes in a pooled sample, and Jayson Talag, Dario Copetti, and the Arizona Genomics Institute (AGI) for sequencing services. This work was funded by USDA-NIFA Postdoctoral Fellowship #2024-67012-43394 to J.A.P. Participation by T.R.C. and A.R.P. was supported by USDA-NIFA Postdoctoral Fellowship #2024-67012-43394; A.R.P. was also supported by the University of Arizona Undergraduate Biology Research Program with funding from the University of Arizona Office of Research & Partnerships. K.M.D. was supported by USDA-NIFA #2023-67013-40169. Mention of trade names or commercial products in this publication is solely for the purpose of providing specific information and does not imply recommendation or endorsement by the United States Department of Agriculture (USDA). USDA is an equal opportunity employer and provider.

## DATA AVAILABILITY

Raw sequencing data has been deposited at NCBI under BioProject PRJNA1377148. The genome assembly and annotation are available at Zenodo (doi.org/10.5281/zenodo.17334877) and at NCBI accession JBSUSI000000000. All code for this project can be found at www.github.com/jessiepelosi/NeoGenome.

